# Mechanism of the Temperature-Dependent Self-Assembly and Polymorphism of Chitin

**DOI:** 10.1101/2023.05.24.542201

**Authors:** Aarion Romany, Gregory F. Payne, Jana Shen

## Abstract

Chitin is the second most abundant natural biopolymer; its crystalline structures have been extensively studied; however, the mechanism of chitin’s self-assembly is unknown. Here we applied all-atom molecular dynamics to study chitin’s self-assembly process at different temperatures. Strikingly, at 278 K, an amorphous aggregate was formed, whereas at 300 K single-sheet and at 323 K both single- and multi-sheet nanofibril regions were formed. The nanofibrils contain antiparallel, parallel or mixed orientation chains, with antiparallel being slightly preferred, recapitulating chitin’s polymorphism observed in nature. The inverse temperature dependence is consistent with the recent experiment. The analysis suggested that the multi-sheet nanofibrils are assembled by stacking the single nanofibril sheets, which are formed through two types of pathways in which hydrophobic collapse either precedes or is concomitant with increasing number of interchain hydrogen bonds and solvent expulsion. Furthermore, the antiparallel and parallel chains are mediated by different interchain hydrogen bonds. The analysis also suggested that the inverse temperature dependence may be attributed to the hydrophobic effect reminiscent of the low critical solution temperature phase behavior. The present study provides a rich, atomic-level view of chitin’s polymorphic self-assembly process, paving the way for the rational design of chitin-derived novel materials.

## Introduction

Following cellulose, chitin is the second most abundant naturally occurring polymer on earth. Owing to the biodegradability and high tensile strength, chitin derivatives have been found in diverse applications, such as tissue engineering, drug carriers, cosmetics, wound dressing, and recently pickering emulsion. ^1–6^ However, to make functional materials from chitin, a challenge is to find effective and environmentally friendly ways to dissolve it in solution, ^7,8^ and once dissolved, another challenge is to achieve controlled fabrication. ^4,9^ Thus, a detailed understanding of chitin’s self-assembly mechanism is important.

Chitin is a homopolymer composed of β-(1-4)-linked N-acetylglucosamine units. Native chitin is a semicrystalline fibril material consisting of crystalline (microfibers) and amorphous regions, where the crystal is composed of a 2–5 nm thick fibril made up of 18–25 chains. ^4^ Chitin’s crystalline structures have been studied by numerous groups using electron microscopy and X-ray diffraction for over 50 years, which revealed three (α, β, and γ) polymorphic forms depending on the natural sources (Fig. 1a and 1b). ^10–18^ The most abundant allomorph is α-chitin, which contains antiparallel chains in the crystalline structure and can be found in hard shells of various crustaceans and cell walls of select fungi ^19^ (Fig. 1a). The second common allomorph is β-chitin, which contains parallel chains and can be often found in soft protein-rich structures such as squid pens and tube worms ^20^ (Fig. 1a). γ-chitin is the rare allomorph, which contains both parallel and antiparallel chains and has been found in the cocoon fibers of beetle and stomach of squid ^13,18^ (Fig. 1a). α-chitin is less susceptible to intra-crytalline swelling and is always obtained in recrystallization from solution, which has been attributed to the presence of inter-sheet hydrogen bonds (h-bonds). ^13,19^ Interestingly, although having both chain orientations, γ-chitin is more similar to α-than β-chitin in the X-ray diffraction pattern and thermal stability. ^13^ The mechanism by which biosynthesis leads to the distinct chitin allomorphs is unknown.

**Figure 1.**
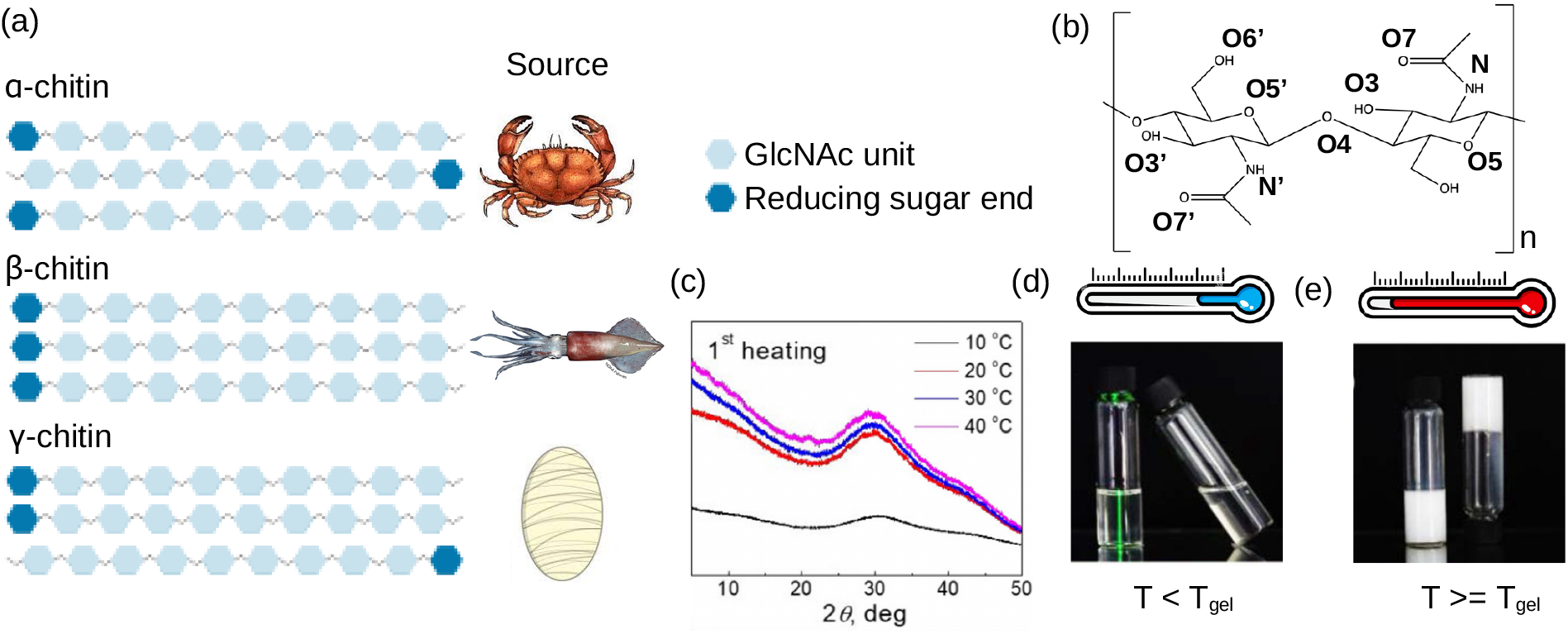
Schematics of chitin’s polymorphism and heat-induced gelation. **(a)** Three allomorphs of chitin and their common natural sources. ^10–18^ The chain direction is illustrated by the reducing sugar end (blue). **(b)** Chemical structure of *β*(1,4)-linked acetylglucosamine (GlcNAc) dimer, the building block of chitin. **(c)** Powder X-ray diffraction pattern of chitin solution (10°C, dissolved in aqueous KOH/urea) and heat-induced gels in the first heating process. **(d)** Below the gelation temperature (T_gel_) chitin solution remains transparent. **(e)** Above T_gel_ chitin solution turned to white solid gel. Images in (c,d,e) are taken from Ref. ^21^ and reproduced with permission from Giant.

Two recent studies of Cai and workers ^21,22^ based on a variety of experimental techniques including atomic force microscopy (AFM), fluorescence microscopy, and wide-angle X-ray diffraction examined the gelation behavior of α- and β-chitin at different concentrations, temperatures, and aging times. Surprisingly, they found that gelation of α-chitin that was dissolved in KOH/urea is heat induced – the diffraction curve of the chitin solution is smooth with minimum intensity at 10°C but the intensity increases dramatically with temperature between 20°C and 40°C ^21^ (Fig. 1c and 1d). A similar behavior was observed for β-chitin which was dissolved in KOH/urea, although the self-assembled hydrogels were made up of α-chitin nanofibrils. ^22^ These observations seem to contradict the common belief that hydrogen bonding, which is thought to be the driving force for the association of chitin chains, ^19^ weakens with increasing temperature. To provide an explanation, the authors hypothesized that the heat-induced gelation is due to increased hydrophobic association of chitin chains at increasing temperature. ^21,22^

Compared to a large body of experimental work on chitin’s physicochemical properties, theoretical investigations have been sparse. A few studies based on molecular dynamics (MD) simulations starting from a prebuilt model crystallite structure of α-chitin examined the decrystallization thermodynamics, ^23^ hydrogen bonding pattern in the nanofiber, ^24^ or binding of a short chitosan oligomer. ^25^ Based on the pre-built α- and β-nanofibers, a recent reactive MD study investigated the different mechanical properties of the two crystalline forms. ^26^ Distinct from the published computational work, we performed MD simulations of chitin self-assembly from solution at different temperatures to understand chitin’s polymorphism and heat-induced self assembly. Strikingly, our simulations captured the formation of nanofibril in α, β and γ allomorphs, and in agreement with the experiments of Cai and coworker, ^21,22^ the self-assembly did not occur at 278 K (5°C), while the nanofibril formation is more ordered at 323 K (50°C) as compared to 300 K (27°C). The analysis revealed the temperature-dependent conformational dynamics and self-assembly process of the chitin chains. Furthermore, the relative orientations of the chitin chains in the nanofibers were found to be attributed to the distinct interchain hydrogen bonds (h-bonds). Together, our data provides a rich and intimate view of the temperature-dependent polymorphic self-assembly process of chitin.

## Results and Discussion

### Conformational dynamics of the dissolved chitin chains is slightly enhanced at 323 K

The self-assembly simulations of chitin were initiated from 24 chains, each comprised of 10 N-acetyl-glucosamine units randomly distributed in a 85 Å x 85 Å x 85 Å cubic box filled with water and periodically replicated in three dimensions. The simulation was performed at three temperatures, 278 K, 300 K, and 323 K, and for each temperature, three independent simulations were initiated from different random velocity seeds. Each simulation was continued until the total solvent accessible surface and the total number of inter-chain hydrogen bonds plateaued, for 3 μs at 278 K and 2 μs at 300 and 323 K (Fig. S1 and S2).

We first investigated if the conformation and/or dynamics of the dissolved chitin chains are affected by temperature. To do so, we calculated the Φ and Ψ dihedral angles around the backbone glycosidic bond (Fig. 2a) of the chitin chains before the self-assembly process began (the initial 50 ns data was used). Following the convention used in the previous work, ^27,28^ Φ and Ψ torsion angles are defined as H1^*′*^-C1^*′*^-O4-C4 and H4-C4-O4-C1^*′*^, respectively (Fig. 2a). The free energy surface (FES) as a function of Φ and Ψ (Fig. 2b) displays a global minimum near (45^*°*^, 0^*°*^) for all three temperatures, suggesting that the dominant conformation the chitin chains assume is *syn*, which is in agreement with the recent NMR and simulation data for β(1 → 4)-linked disaccharides in aqueous solution ^28^ as well as our previous simulations of chitosan chains in solution. ^27^

**Figure 2.**
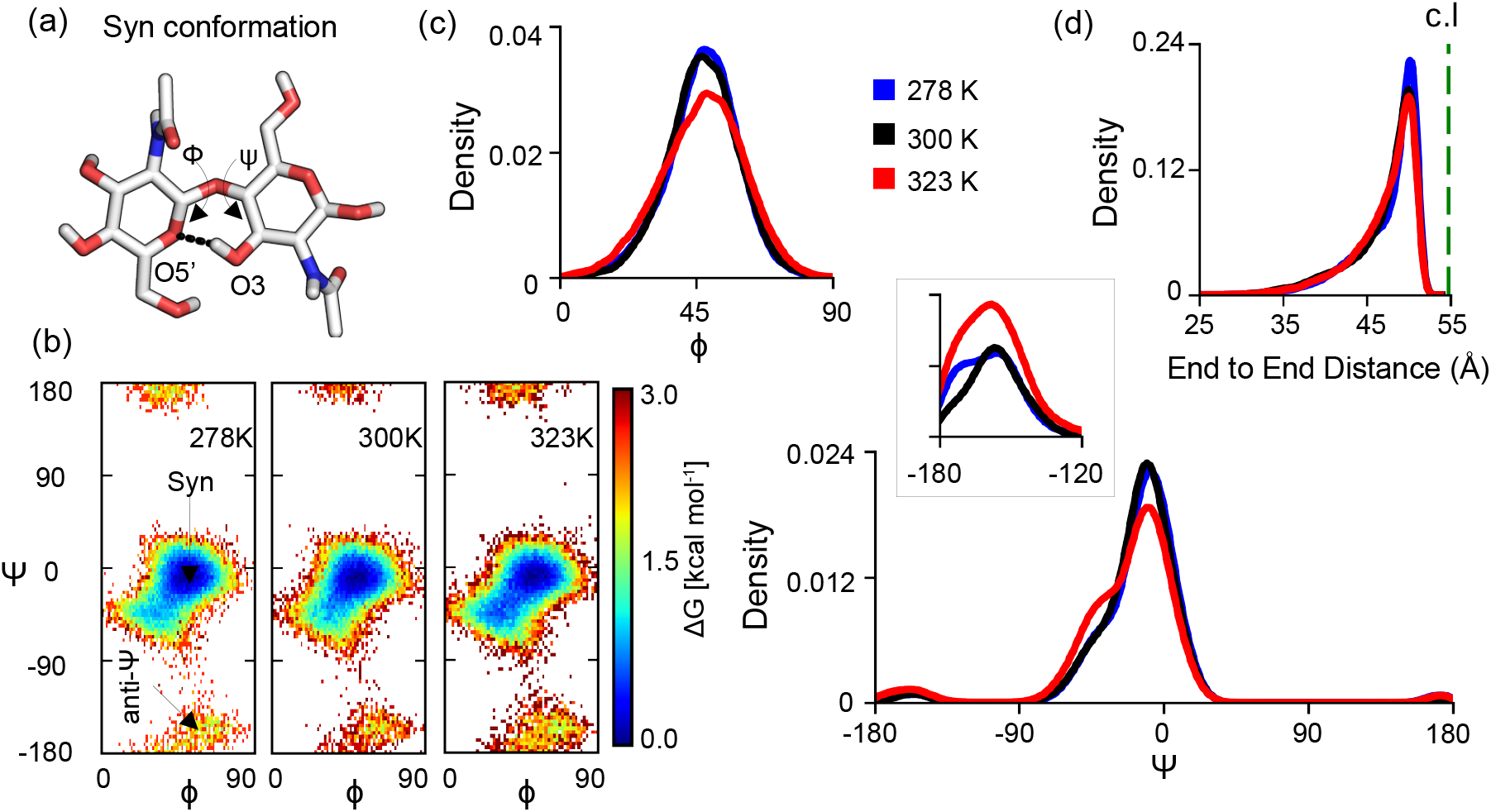
Conformational dynamics of the dissolved 10-mer chitin chains slightly enhanced at 323 K. **(a)** Structure of an acetylglucosamine dimer in a *syn* conformation. The glycosidic backbone Φ and Ψ angles as well as the O3H-O5^*′*^ intramolecular h-bond are indicated. In the alternative anti-Ψ conformation, the sugar rings are perpendicular to each other. ^27^ **(b)** Free energy surface (FES) as a function of the Φ and Ψ angles for the dissolved chitin chains at three temperatures. The *syn* and anti-Ψ regions are indicated. **(c)** Probability distributions of the Φ (top) and Ψ (bottom) angles of the dissolved chitin chains at 278 (blue), 300 (black), and 323 K (red). **(d)** Probability distributions of the end-to-end distances (between the O1 atom of the first and the O4 atom of the last acetylglucosamine unit in the chitin chain) of the dissolved chains at 278, 300, 323 K. Data for each temperature is the aggregate of the initial 50 ns of the three simulation runs. Any chains that are within 4.5 Å of another were excluded. This was further confirmed by visual inspection. The theoretical contour length is indicated as a green dashed line.

A closer examination of the FES reveals that the probability of the anti-Ψ region is slightly higher at 323 K (Fig. 2b). This is related to the slightly broader distribution of the Φ angle and greater sampling of the alternate Ψ angles at 323 K as compared to 278 and 300 K (Fig. 2c), suggesting that the solution chitin chains have a more flexible backbone at 323 K. To investigate if temperature also affects the dimension of the solution chitin chains, we calculated the distribution of the end-to-end distance (Fig. 2d). The peak of the distribution for 323 K is slightly left shifted and lower compared to 278 and 300 K, demonstrating that the chains are slightly less extended and more flexible at 323 K, which is in accord with the more flexible backbone discussed earlier (Fig. 2b,c).

### Increasing temperature promotes chitin’s nanofibril formation

We next examined the self-assembly process of the chitin chains at three temperatures. At the end of all simulation runs, the chitin chains formed a single aggregate with at most one chain left in solution (see representative final snapshots in Fig. 3a). To examine the degree of order, we first calculated the distributions of the angles between any two chitin chains. At 273 K, the distribution is nearly uniform with many local peaks between 0^*°*^ and 180^*°*^, indicating that the chains are largely randomly oriented (Fig. 3b). In contrast, the distribution at 300 K is bimodal, showing the major mode (sharp peak) near 180^*°*^ and the minor mode (lower and broader peak) near 0^*°*^, which indicates that many chains are antiparallel and some are parallel to each other (Fig. 3b). At 323 K, the bimodal distribution is more sharply peaked; the minor peak near 0^*°*^ has an increased height and decreased width, indicating that the chains are mostly aligned antiparallel/parallel to each other with the antiparallel orientation being slightly favored as the 300 K distribution (Fig. 3b). This is consistent with the experimental observation that α-chitin which contains all antiparallel chains is the most common and stable allomorph. ^13,19^

**Figure 3.**
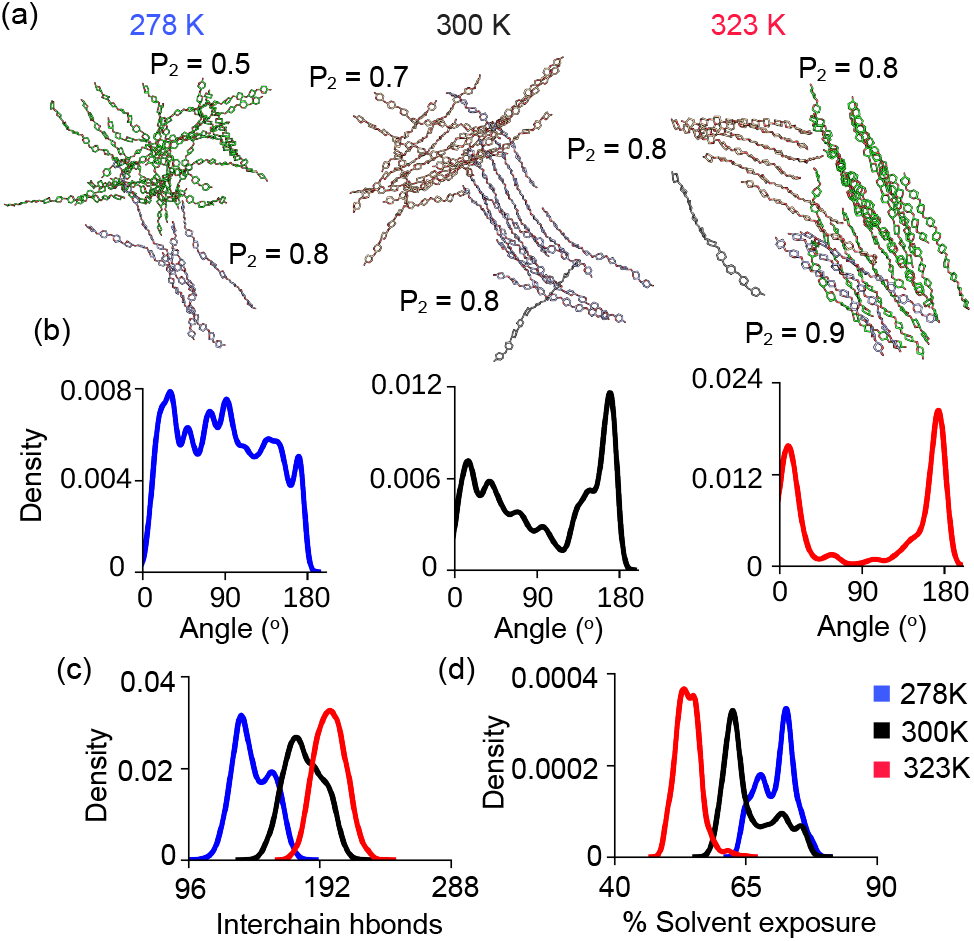
Temperature effect on the self-assembled structure, hydrogen bonding and solvent exclusion between chitin chains. **(a)** Representative final snapshots from the simulations at three temperature showing the formed aggregate and nanofibril regions. Only the sugar rings and glycosidic linkages are shown for clarity. Snapshots at 278 K, 300 K and 323 K are taken from run 3, 3 and 2, respectively. Ordered regions (shown in different colors) are identified in the aggregates, with the local *P*_2_ order parameter given. *P*_2_ *>*0.7 indicates a mostly ordered state. The final 100 ns of simulation time was used for the *P*_2_ calculation. **(b)** Probability distribution of the angle between chains at three temperatures. **(c)** Probability distribution of the total number of interchain h-bonds for the 24 chitin chains at three temperatures. **(d)** Probability distribution of the percentage solvent exposure of the 24 chitin chains at three temperatures. Solvent exposure was calculated as the solvent accessible surface area (SASA) of the chitin heavy atoms relative to the value calculated following the energy minimization of the system. For (b), (c) and (d), the final 100 ns of three trajectories were combined for each temperature.

To quantify the degree of order, we calculated a P_2_ order parameter analogous to the global order parameter used for characterizing nematic liquid crystals and amyloid nanofibrillation ^29^ (see Methods). P_2_ is bound between 0 and 1, where 0 indicates a completely disordered state with the chains randomly oriented relative to each other, and 1 indicates a perfect fibrillar state in which the chains are parallel or antiparallel to each other. Following our previous work on amyloid fibrillation, ^29^ we consider the chitin chains as in a fibrillar or an amorphous state if P_2_ is >0.7 or < 0.3, respectively. The P_2_ values for the aggregates formed in the three runs at 278 K, 300 K and 323 K are 0.4/0.4/0.5, 0.3/0.2/0.6, 0.8/0.8/0.4, respectively (Table S1).

Visualization of the aggregates suggested that the chitin chains can be grouped into two or more regions and some of the regions are remarkably ordered, resembling nanofibrils (Fig. 3a). Thus, we calculated the order parameter for each region (Table S1). At 278 K, two regions, one comprised of 4 or 5 chains and another one comprised of 19 or 20 chains, were identified in the three runs. While the first region is somewhat ordered (P_2_ of 0.7 or 0.8 in two runs), the second region is closed to being amorphous (P_2_ of 0.4 or 0.5) in all three runs (Fig. 3a and Table S1). At 300 K, 2–5 regions were identified in the three runs, and most of them were more ordered than those formed at 278 K (Fig. 3a and Table S1). Run 1 resulted in three highly ordered regions comprised of 3 or 7 chains (P_2_ of 0.9 or 1.0), one somewhat ordered region (P_2_ of 0.7), and one amorphous region. Run 2 resulted in one highly ordered region of 5 chains (P_2_ of 0.9) and the remaining ones are amorphous. Run 3 resulted in two ordered regions comprised of 10 and 13 chains (P_2_ of 0.7 and 0.8). At 323 K, all three runs showed highly ordered regions with P_2_ of 0.8 or greater. The largest region contains 21 chains with a P_2_ value of 0.8, followed by the regions with 14 and 10 chains and respective P_2_ values of 0.8 and 0.9 (Fig. 3a and Table S1). Comparing the data across the three temperature, it is evident that larger, more ordered (i.e., fibril-like) regions are formed with increasing temperature. In the discussion that follows, we will refer to the regions with P_2_>0.7 as nanofibrils except for those from the 278 K simulations which lack of specific interchain h-bonds (see later discussion).

### Interchain hydrogen bond formation increases and solvent accessibility decreases with increasing temperature

To examine the effect of temperature on chitin’s self-assembly, we also calculated the probability distributions of the total number of interchain h-bonds and the percentage solvent exposure of the 24 chitin chains based on the last 100 ns of the simulations at three temperatures. The peak of the distribution of the interchain h-bond number is shifted towards larger values as temperature increases from 278 K to 323 K (Fig. 3c), indicating that more interchain h-bonds are formed with higher temperature, which is consistent with the increased order of the formed aggregate (Fig. 3a). Although this relationship between temperature and hydrogen bonding may seem contrary to the common belief, it is in agreement with the experimental observation that the sol-gel transition of dissolved chitin occurs above, rather than below, a gelation temperature (around 288 K for dissolved α-chitin). ^21,22^

Next, we examined the percentage solvent exposure of the 24 chitin chains, which is calculated as their solvent accessible surface area (SASA) relative to the fully-solvent exposed state. As temperature increases, the distribution is shifted towards smaller values (Fig. 3d), indicating that the chitin chains become more buried or in other words, more water is displaced from the first solvation shells. This suggests that the temperature-dependent self-assembly of chitin may be explained by the hydrophobic effect. In the classical view, the association of small hydrocarbons in water is driven by the hydrophobic effect (i.e., -TΔS) which becomes more negative with increased temperature up to a certain temperature. ^30,31^ In a recent MD study; however, Durell and Ben-Naim found that between 280 and 360 K the free energy of bringing together hydrophilic solutes that are directly h-bonded to each other also becomes more negative with increasing temperature as does for two hydrophobic solutes. ^32^ Their analysis ^32^ demonstrated that the increase in the entropic term TΔS is due to the release of h-bonded water as the solutes form h-bonds among themselves, which may explain the increased interchain h-bonds and displacement of the first-solvation-shell (i.e., bound) water for the temperature-dependent self-assembly of chitin. Given the high correlation between interchain h-bonding and solvent accessibility, we begin to see the important role solvent plays in the self-assembly of chitin.

### The formation of single-sheet nanofibrils proceeds via two distinct mechanisms

In theoretical studies of protein folding pathways, one often considers a low-dimensional free energy surface (FES) spanned by order parameters such as the fraction of native contacts (ρ) and the radius of gyration (R_*g*_). Based on the shape of the FES spanned by ρ and R_*g*_, two types of mechanisms have been proposed for protein folding. ^33^ In the first mechanism which is represented by a L-shaped FES, ^33^ protein collapses in solvent first before rearrangement to increase the native contacts. In the second mechanism which is represented by a diagonally-shaped FES, ^33^ the hydrophobic collapse and native contact formation are concomitant. It is thought that the proteins with a great number of long-range contacts (e.g. β-sheets) follow the L-shaped profile, whereas proteins with predominantly short-range contacts (e.g. helices) follow the diagonal profile. ^33^ Since chitin fibrils are mainly stabilized by long-range interchain h-bonds, we hypothesized that the self-assembly pathways follow a L-shaped profile.

To test the above hypothesis, we calculated the FES for the formation of individual nanofibril regions listed in Table S1. As order parameters, we used R_*g*_, the number of interchain h-bonds per GlcNAc unit (in analogous to ρ for protein folding), and the percentage solvent exposure (based on solvent accessible surface area or SASA calculations). Interestingly, both L- and diagonal-shaped FES’ were observed for a single-sheet nanofibril formation. Two examples are given in Fig. 4. In the first example, the formation of a 6-chain nanofibril sheet follows a L-shaped FES, indicating that in the first stage of the self-assembly the chitin chains move closer together while the number of interchain h-bonds remains minimal (snapshot 1 and 2 in Fig. 4a and FES in Fig. 4b left). In this stage, the percentage solvent exposure (which measures the water bound to the chitin chains) of the chitin chains does not significantly change (see FES in Fig. 4b right), indicating that while the bulk water is displaced the chitin chains remain bound to water, i.e., the first solvation shell is unperturbed. Consistent with the minimal number of interchain h-bonds, the chains are randomly orientated towards each other (snapshot 1 and 2 in Fig. 4a). Following the chain collapse stage, rearrangement of the chains occurs (snapshot 3 and 4 in Fig. 4a), which appears to be driven by an increase in the number of interchain h-bonds while the R_*g*_ remains more or less constant (Fig. 4b left). At the same time, the percentage solvent exposure of the chains decreases from over 80% to less than 55% (Fig. 4b right), indicating that the bound water are expelled as more h-bonds are formed between the chitin chains. With the increase in the number of h-bonds, the chains become more aligned with each other as evident from the increase in the P_2_ value. Snapshot 4 shows a 6-chain single-sheet nanofibril (P_2_ of 0.9) with most chains perfectly aligned (Fig. 4a).

**Figure 4.**
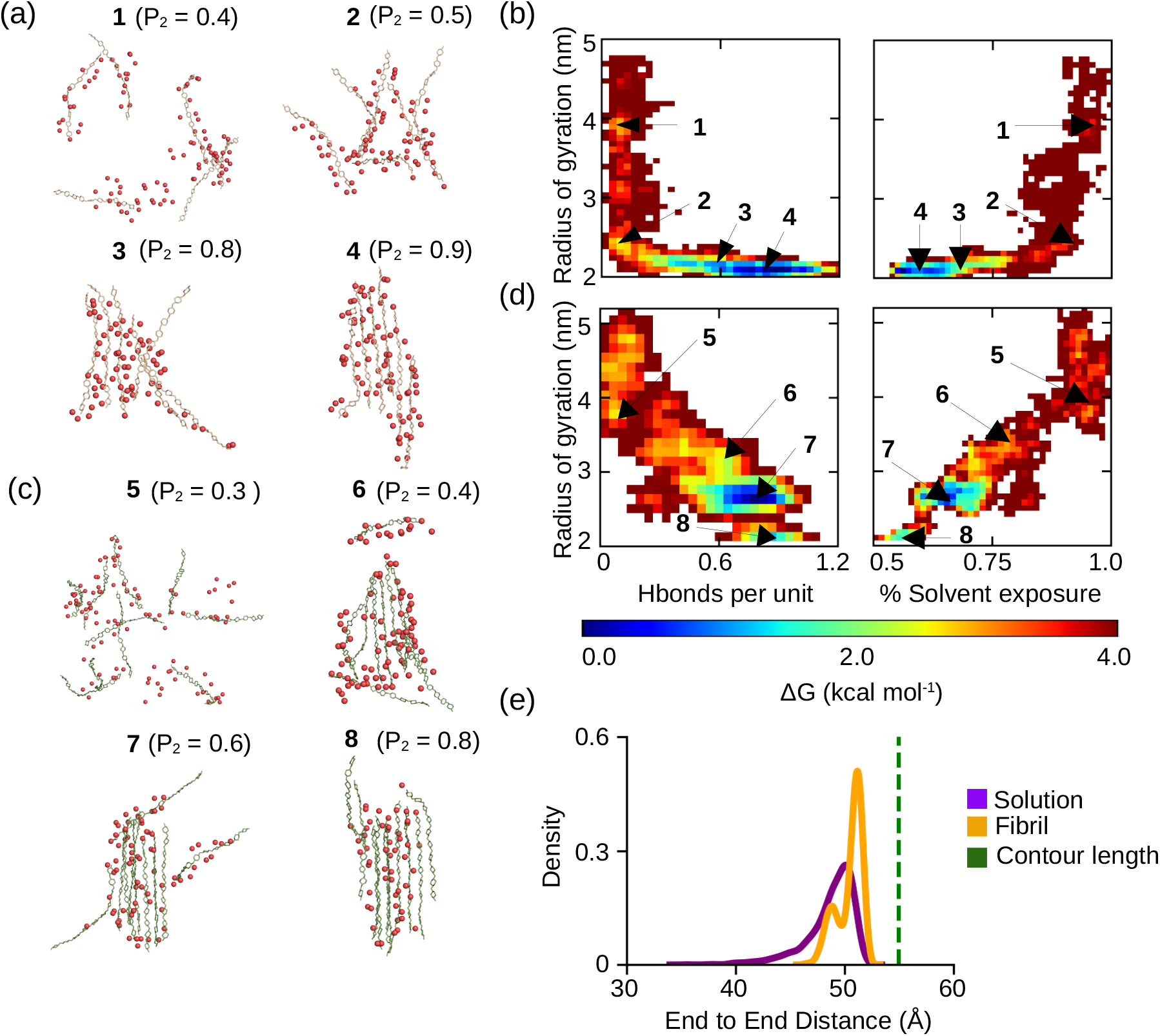
The self-assembly pathway of a single-sheet nanofibril of chitin follows two mechanisms. **(a, c)** Representative snapshots extracted from the local minimum regions in the FES to illustrate the formation of a 6-chain (a, snapshot 1–4) and a 10-chain (c, snapshot 5–8) single-sheet nanofibrils. Red spheres represent water molecules in the first hydration shell defined as within a 3.4-Å radius from any heavy atom of the chain. **(b, d)** FES spanned by the radius of gyration and the number of h-bonds per GlcNAc unit (left panel) or the percentage solvent exposure (right panel) of the chitin chains. The number of h-bonds per unit was calculated by dividing the total number of interchain h-bonds by the number of chains and the degree of polymerization per chain (the two end units were ignored). The percentage of solvent exposure was calculated using SASA (see Fig. 3). **(e)** The distribution of the end-to-end distance of the chitin chain in solution (purple) and fibril (brown) state. The green dashed line represents the theoretical contour length of 55 Å for a 10-mer chitin chain. Data for (a, b, e) and (c, d) refer to the chains in the nanofibril region 1 formed in the 323 K simulation run 2 and 3, respectively.

In the second example, the formation of a 10-chain single-sheet nanofibril follows predominantly a diagonal-shaped FES (Fig. 4c and 4d). Although there is an initial descent in the R_*g*_ down to the local minimum region 5 (snapshot 5 in Fig. 4c) similar to the transition 1 → 2 for the 6-chain fibril, the transition 5 → 6 → 7 → 8 follows a diagonal FES, indicating that as the chains come together the interchain h-bonds form concomitantly (snapshot 5 in Fig. 4d, left). This transition is also accompanied by a steep decrease in the percentage solvent exposure, indicating that the first-solvation shell water is expelled from the chains. These data suggest that unlike protein folding, both types of pathways are possible for chitin’s nanofibrillation.

To test if there is a change in the chitin chain dimension due to fibrillation, we plotted the probability distribution of the end-to-end distance of the chains in solution (first 50 ns data) and in the nanofibril (final 50 ns data) using the chains that formed a nanofibril sheet at 323 K (data used for Fig. 4a and 4b). The chitin chains in solution (before the self-assembly began) show a broader and slightly left-shifted distribution as compared to the chains in the fibrillar state (Fig. 4e). This suggests that the chains are more flexible in solution and they become more extended and rigid (rodlike) when transitioning to the fibrillar state, which is likely a result of restricted motion due to interchain h-bonding and van der Waals interactions.

The assembly process of multi-sheet nanofibrils appears to have two stages (two examples are given in Fig. 5). In the first example, three nanofibril sheets are formed first by following a predominantly diagonal-shaped FES with concomitant chain collapse, interchain h-bond formation and water expulsion (1 → 2 → 3, Fig. 5a and 5b). In the second stage, the three fibril sheets move closer to form stacking by a small decrease in the solvent exposure (some water in the first solvation shell is expelled), and interestingly, one chain from the solution is added onto one sheet which provides further stabilization (3 → 4, Fig. 5a and 5b). In the second example, two nanofibril sheets are formed first by following a L-shaped FES, (5 → 6 → 7, Fig. 5c and 5d). In the second stage, the two sheets move closer to form stacking followed by taking up one chain from solution (7 → 8 → 9, Fig. 5c and 5d). These data suggest that the individual fibril sheets are formed first before inter-sheet stacking occurs. As a solution chitin chain is added onto a stacked sheet in both examples, we hypothesize that a multi-sheet nanofibril may serve as a nucleus for chitin’s fibril growth similar to amyloid fibrils. ^34^

**Figure 5.**
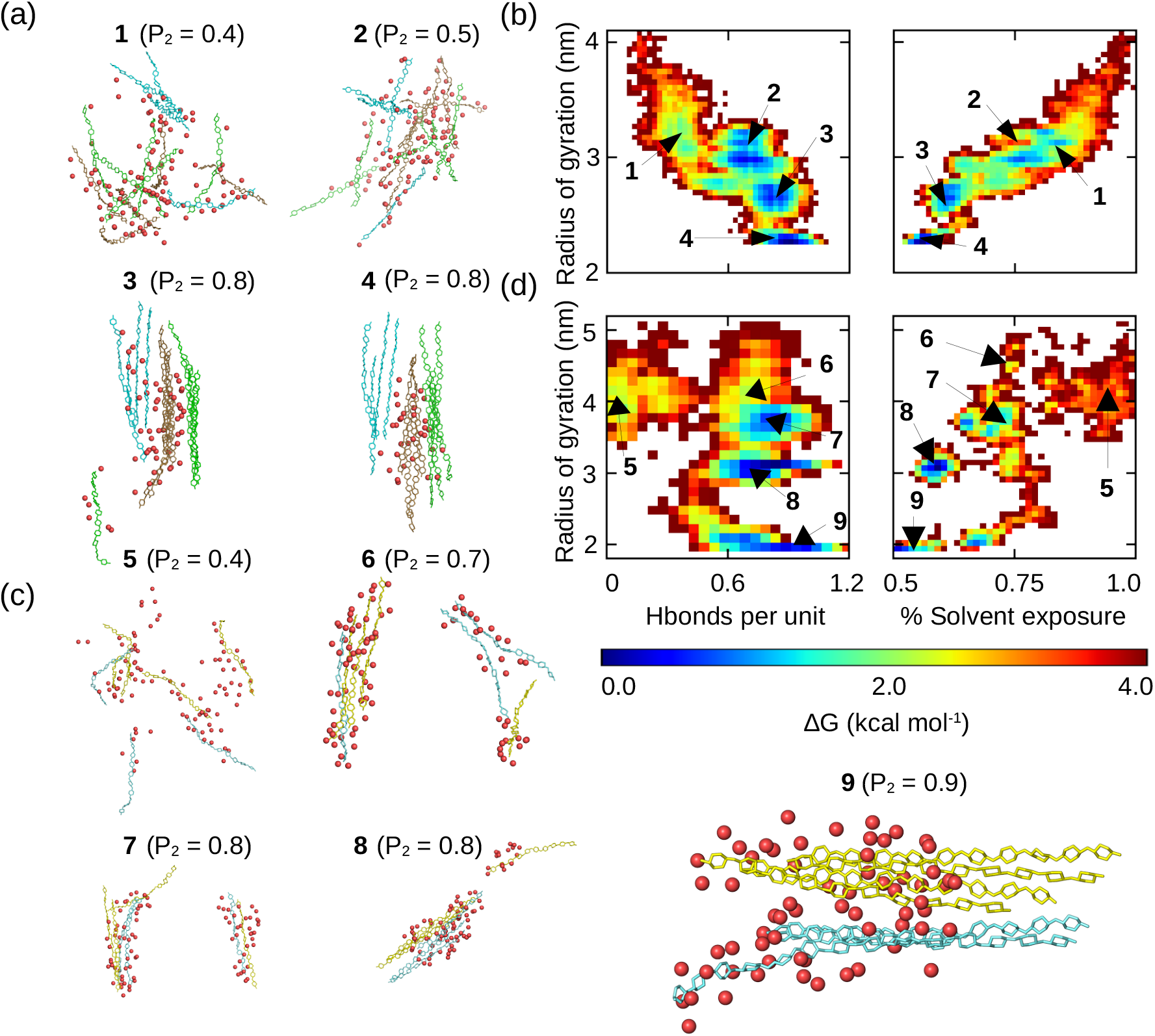
A multi-sheet nanofibril is assembled by stacking the single-sheet nanofibrils. **(a, c)** Representative snapshots extracted from the local minimum regions of the FES to illustrate the formation of a 14-chain three-sheet (a, snapshot 1–4) and a 7-chain two-sheet nanofibril (c, snapshot 5–9). Different sheets are indicated by different colors. **(b, d)** FES spanned by the radius of gyration and the number of interchain h-bonds per GlcNAc unit (left panel) or the percentage solvent exposure (right panel) of the chitin chains. Data for (a, b) and (c, d) refer to the chains in the nanofibril region 3 formed in the 323 K simulation run 2 and 3, respectively.

### Self-assembled nanofibrils are polymorphic

In nature, chitin can self-assemble into three allomorphs distinguished by the chain alignment, parallel (β), antiparallel (α), and mixed (γ). ^10–18^ Interestingly, all three allomorphs are observed in the nanofibrils identified in the final “aggregates” (Table S1), which is consistent with the angle analysis showing both parallel or antiparallel chain orientations at 300 K and 323 K (Fig. 3b). At 300 K, one nanofibril (3 chains) shows parallel, three nanofibrils (3 or 5 chains each) show antiparallel, and two nanofibrils (10 and 13 chains) show mixed alignment. At 323 K, one nanofibril (3 chains) shows parallel, two nanofibrils (3 chains each) show antiparallel, and 6 nanofibrils (3, 14, 21, 10, 7 chains) show mixed alignment. These data suggest that while the smallest nanofibrils can adopt exclusively parallel or antiparallel orientations, larger nanofibrils tend to show mixed chain alignment.

### Specific interchain hydrogen bonds mediate fibril formation and polymorphism

What forces give rise to the single- and multi-sheet nanofibrils and the different allomorphs? Interestingly, only the transient O6H…O7 h-bonds were found the nanofibril sheets (Fig. S3); instead, the sheet stacking was mainly mediated by van der Waals contacts, suggesting that hydrophobic effect is dominant for inter-sheet stabilization (Fig. 6a). This is in consistent with the experiments. The X-ray diffraction data of β-chitin demonstrated an absence of h-bonds between the fibril sheets, ^16^ while a Fourier transform infrared (FTIR) study suggested that the α-chitin may contain O6^*′*^H…O7 h-bonds. ^14^

**Figure 6.**
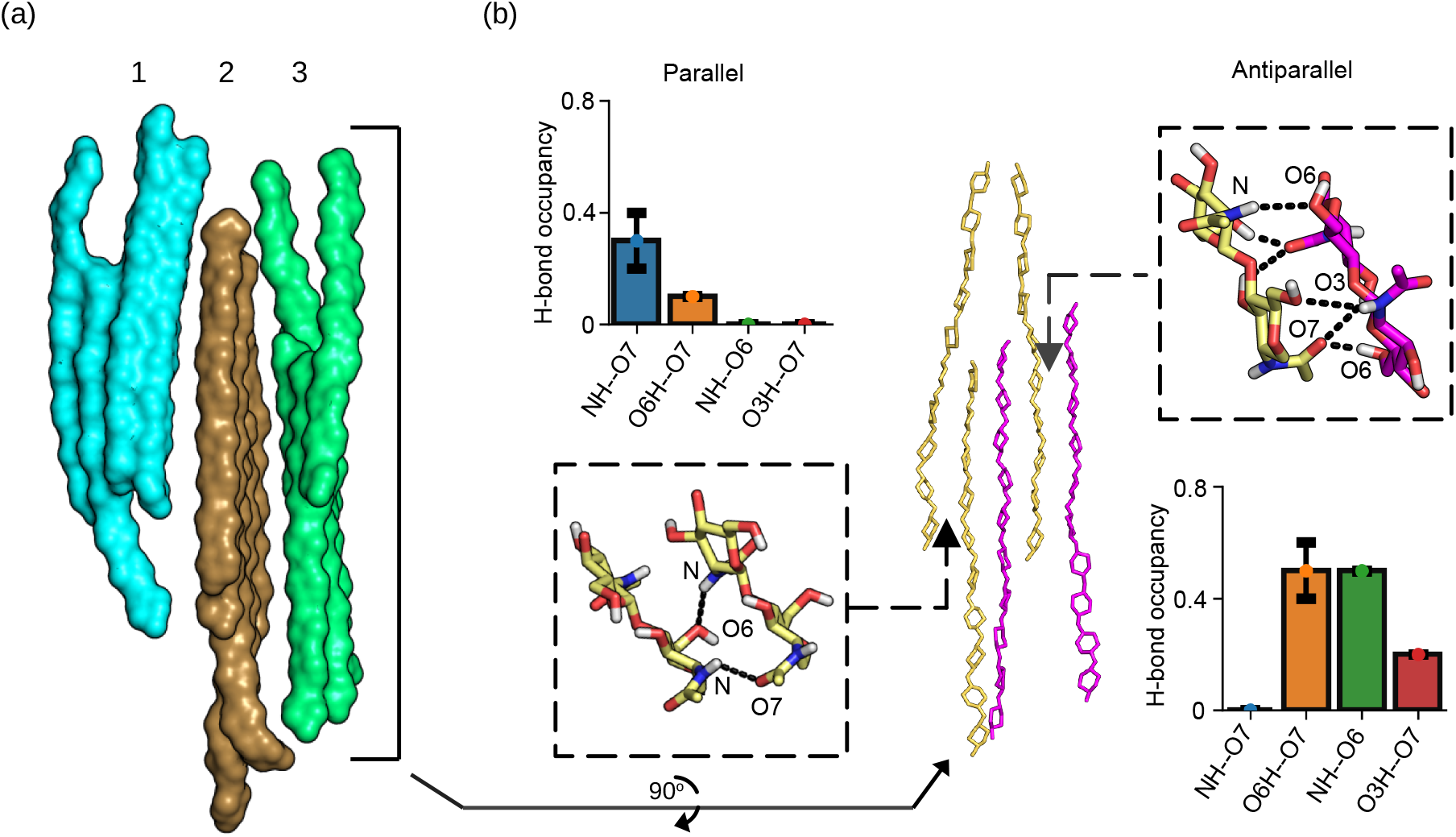
Characterization of the chain orientations and interchain h-bonding in the chitin nanofibril. **(a)** A snapshot of an example nanofibril (in the van der Waals surface rendering) comprised of three sheets, and each sheet has 4–5 chains. Data from the 323 K simulation run 2 is used. **(b)** A zoomed-in view of sheet 3 (green in a) showing the intra-sheet chain orientations and h-bonding. Brown and magenta colors indicate the two chain directions. The occupancies and the zoomed-in views of the specific h-bonds that mediate the parallel and antiparallel chains are given on the left and right panels, respectively. In the zoomed-in view, only two GlcNAc units are shown. The occupancies were calculated from all nanofibril sheets in each of the 323 K runs. The average and standard deviation over the three runs are shown. The final 100 ns of the trajectories was used.

Turning to the intrasheet interactions, we examined specific interchain h-bonds. Surprisingly, despite a large number of interchain h-bonds at 278 K (Fig. 3b), the occupancies of specific h-bonds are very low (the most occupied h-bond O6H…O7 is below 10%, Fig. S4), which explains the increased disorder and lack of fibril formation (Fig. 3a). By contrast, in the simulations at 300 K and at 323 K, four specific interchain h-bonds, NH… O7, O6H… O7, NH… O6, and O3H… O7, were found, and interestingly, parallel and antiparallel chains are stabilized by different h-bonds (Fig. 6b and Fig. S5). The parallel configurations are stabilized by the NH… O7 h-bond with an occupancy of about 30% (averaged across the three runs at 323 K) and the O6H… O7 h-bond with an occupancy of 10% (Fig. 6b, left panel). In contrast, the antiparallel configurations are stabilized by the O6H… O7 (average occupancy ∼50%), NH… O7 (average occupancy ∼50%), and and O3H… O7 (average occupancy ∼20%) h-bonds (Fig. 6b, right panel). Overall, the antiparallel configurations have more stable interchain h-bonds than the parallel configurations, which provides an explanation for the increased thermal stability ^12^ and decreased susceptibility to solvent-mediated swelling ^19^ of α-chitin which is formed by antiparallel chains relative to β-chitin which is formed by parallel chains.

## Conclusions

The MD simulations offered an atomic-level description of how chitin chains self assemble into nanofibrils from solution at three temperatures. At 278 K a largely amorphous aggregate without specific interchain h-bonds was formed, whereas at 300 and 373 K nanofibril regions of various sizes were identified within the final aggregates. At 300 K, single-sheet nanofibrils comprised of 3-13 chains were assembled; while at 323 K, nanofibrils with two or three sheets with up to 21 chains in total were assembled. These data are consistent with the recent observation of the heat-induced gelation of dissolved chitin. ^21,22^ The analysis showed that as temperature increases from 278 to 323 K, the total number of intermolecular h-bonds increases and more solvent is expelled from the first-solvation shells of the chitin chains. Further analysis revealed that the single-sheet nanofibril formation of chitin follows two types of pathways: a L-shaped FES profile in which a hydrophobic collapse is followed by maximizing the number of interchain h-bonds while expelling solvent; and the diagonal-shaped FES profile in which the hydrophobic collapse and h-bond formation/solvent expulsion are concomitant. This is in contrast to proteins, which are believed to prefer the L-shaped mechanism when long-range native contacts (e.g., β-sheets) are dominant. ^33^ Importantly, the multi-sheet nanofibrils are assembled by stacking the single sheets largely via van der Waals interactions while further excluding solvent.

Considering the correlation between the interchain h-bond formation and water expulsion (release of h-bonded water) and the hydrophobic nature of intersheet stacking, the inverse temperature dependence of chitin’s self-assembly may be explained by the hydrophobic effect (-TΔS) which dominates over the enthalpic contribution (ΔH). This is consistent with a recent study by Durell and Ben-Naim which found that increasing temperature is more favorable for bringing two hydrophilic solutes together that can form direct h-bonds. ^32^ The inverse temperature dependence of chitin’s self-assembly is reminiscent of the lower critical solution temperature (LCST) behavior, for example of elastin-like peptide, which also contains both hydrophobic and hydrophilic residues and undergoes phase transition into peptide-rich and water phase upon heating. ^35^

The nanofibrils formed in the simulations captured all three naturally occurring allomorphs of chitin: with all parallel, all antiparallel, or mixed chain orientations. Interestingly, the antiparallel orientation was found to be slightly favored over the parallel orientation. Analysis showed that the different orientations are mediated by different h-bonds with the antiparallel chains forming a larger number of h-bonds, which provides an explanation to the experimental observation that α-chitin (formed only by antiparallel chains) is the most stable allomorph. ^13,19^ Our data is consistent with the view that the formation of the specific allomorph (α, β or γ) is due to the microenvironment (e.g., protein wrapping in squid pen) which favors one chain orientation vs. the other. Taken together, the present study provides a rich, atomic-level view of chitin’s self-assembly, which may assist the design of controlled fabrication of chitin-based materials.

## Methods and Protocols

All-atom molecular dynamics simulations were performed using the AMBER20 program. ^36^ The chitin chains were represented by the CHARMM36 carbohydrate force field. ^37,38^ 24 chains, each composed of 10 acetylglucosamine units were randomly distributed using PACKMOL ^39^ in a 85 Å x 85 Å x 85 Å cubic box filled with CHARMM-style TIP3P water model. ^40^ The simulation system comprised approximately 57,978 atoms, and the weight percentage of the chitin was 17.2 wt%, which is about three times the experimental concentration of 6 wt%. ^21,22^ The higher concentration was used to accelerate the self-assembly, and is not expected to affect the mechanisms investigated here. CHAMBER ^41^ was used to convert the CHARMM parameter and topology files to AMBER format.

A total of 9 simulations were conducted of the system. Three independent simulations were performed for each system at each temperature of the three temperatures (278K, 300K or 323K) starting from different random velocity seeds. The system first underwent energy minimization with a harmonic restraint potential (force constant of 10 kcal/mol/Å ^2^) placed on the heavy atom positions. The system was then heated over 1 ns to one of the three temperatures in the NVT ensemble. The system was then equilibrated for a total of 100 ns in which the harmonic potential force constant was gradually reduced from 5, 2.5, 1, 0.1 to 0 kcal/mol/Å ^2^ under constant NPT conditions. The temperature was maintained using a Langevin thermostat, ^42^ and the pressure was maintained at 1 bar using the the Monte Carlo barostat. ^43^ Each production run lasted between 2 and 3 μs under the constant NPT conditions. The van der Waals interactions were smoothly switched to zero from 10 to 12 Å. The particle mesh Ewald method ^44^ was used to calculate long-range electrostatic energies with a sixth-order interpolation and 1 Å grid spacing. Bonds involving hydrogens were constrained using the SHAKE algorithm ^45^ to enable a 2-fs time steps. Subsequent analysis of simulations was done using cpptraj ^46^ and VMD. ^47^

The relative chain orientation was calculated as:

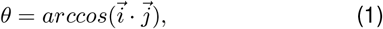

Where 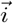 is the molecular vector, connecting the center of masses of the second and ninth sugar rings in an individual chain. 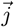 is the vector of the nearest neighbor chain of 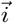. If θ 0^*°*^, then chains are aligned parallel. If θ 180^*°*^, chains are aligned antiparallel. The P_2_ order parameter was defined as^29^

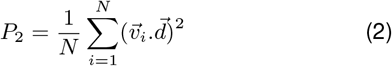

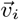 is the normalized molecular vector connecting the center of mass of the second and ninth sugar rings in an individual chain. 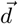 is the director, which is a normalized average vector of the unsigned orientations of all individual 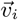. N is the number of molecular vectors (number of chains).

## Supporting Information

A supplemental Table contains the detailed information about the nanofibril regions in the simulation runs. Supplemental figures contain additional analysis of the data.

## Acknowledgements

Financial support from the National Science Foundation (CBET1932963) is acknowledged.

