## Supplemental Information for "Mechanism of the Temperature-Dependent Self-Assembly and Polymorphism of Chitin"

### List of Tables

S1 Composition, order, and chain alignment of the local regions identified in the final aggregate formed in the simulation runs$^a$ . . . . . . . . . . . . . . . . S-3

### List of Figures

S1 Time series of the interchain hydrogen bond formation of chitin chains . . . S-4
S2 Time series of the fraction of maximal solvent accessible surface area of the chitin chains . . . . . . . . . . . . . . . . . . . . . . . . . . . . . . . . . . . S-5
S3 Transient intersheet hydrogen bond. . . . . . . . . . . . . . . . . . . . . . . . S-6
S4 Specific Hbonds at low temperature. . . . . . . . . . . . . . . . . . . . . . . . S-6
S5 Specific Hbonds at 300K . . . . . . . . . . . . . . . . . . . . . . . . . . . . . S-7

### Supplemental Tables

Table S1: Composition, order, and chain alignment of the local regions identified in the final aggregate formed in the simulation runs[a]

| Temperature<br>Run | Region 1<br>#chains ($P_2$) | Region 2<br>#chains ($P_2$) | Region 3<br>#chains ($P_2$) | Region 4<br>#chains ($P_2$) | Region 5<br>#chains ($P_2$) |
|---|---|---|---|---|---|
| | | | **278 K** | | |
| Run 1 | 4 (0.7)<br>para | 19 (0.4)<br>n/a | 1 (n/a)<br>n/a | | |
| Run 2 | 4 (0.2)<br>n/a | 20 (0.4)<br>n/a | | | |
| Run 3 | 5 (0.8)<br>para | 19 (0.5)<br>n/a | | | |
| | | | **300 K** | | |
| Run 1 | 3 (1.0)<br>para | 7 (0.7)<br>mixed | 3 (1.0)<br>anti | 5 (0.9)<br>anti | 6 (0.3)<br>n/a |
| Run 2 | 5 (0.3)<br>n/a | 5 (0.9)<br>anti | 4 (0.1)<br>n/a | 10 (0.3)<br>n/a | |
| Run 3 | 10 (0.7)<br>mixed | 13 (0.8)<br>mixed | 1 (n/a)<br>n/a | | |
| | | | **323 K** | | |
| Run 1 | 3 (0.8)<br>anti | 21 (0.8)<br>mixed, 3 sheets | | | |
| Run 2 | 6 (0.8)<br>mixed | 3 (0.9)<br>mixed | 14 (0.8)<br>mixed, 3 sheets | 1 (n/a)<br>n/a | |
| Run 3 | 10 (0.9)<br>mixed | 3 (0.9)<br>anti | 7 (0.9)<br>mixed, 2 sheets | 4 (1.0)<br>para | |

[a]The $P_2$ parameter was calculated using the final 100 ns of simulation time. 1 refers to one chain remaining in solution. n/a indicates not applicable. Alignment refers to the relative orientation of the chains in the local region: parallel (para), antiparallel (anti), and mixed parallel/antiparallel (mixed). Alignment information is only given for the regions with $P_2 \geq 0.7$. Note, at 278 K, even though region 1 has 5 parallel chains, the occupancies of the specific interchain h-bonds (see Fig. 5 in the main text) are below 10%. Therefore, we do not consider it as a fibrillar state. Multisheet nanofibrils are underlined.

#### Supplemental figures

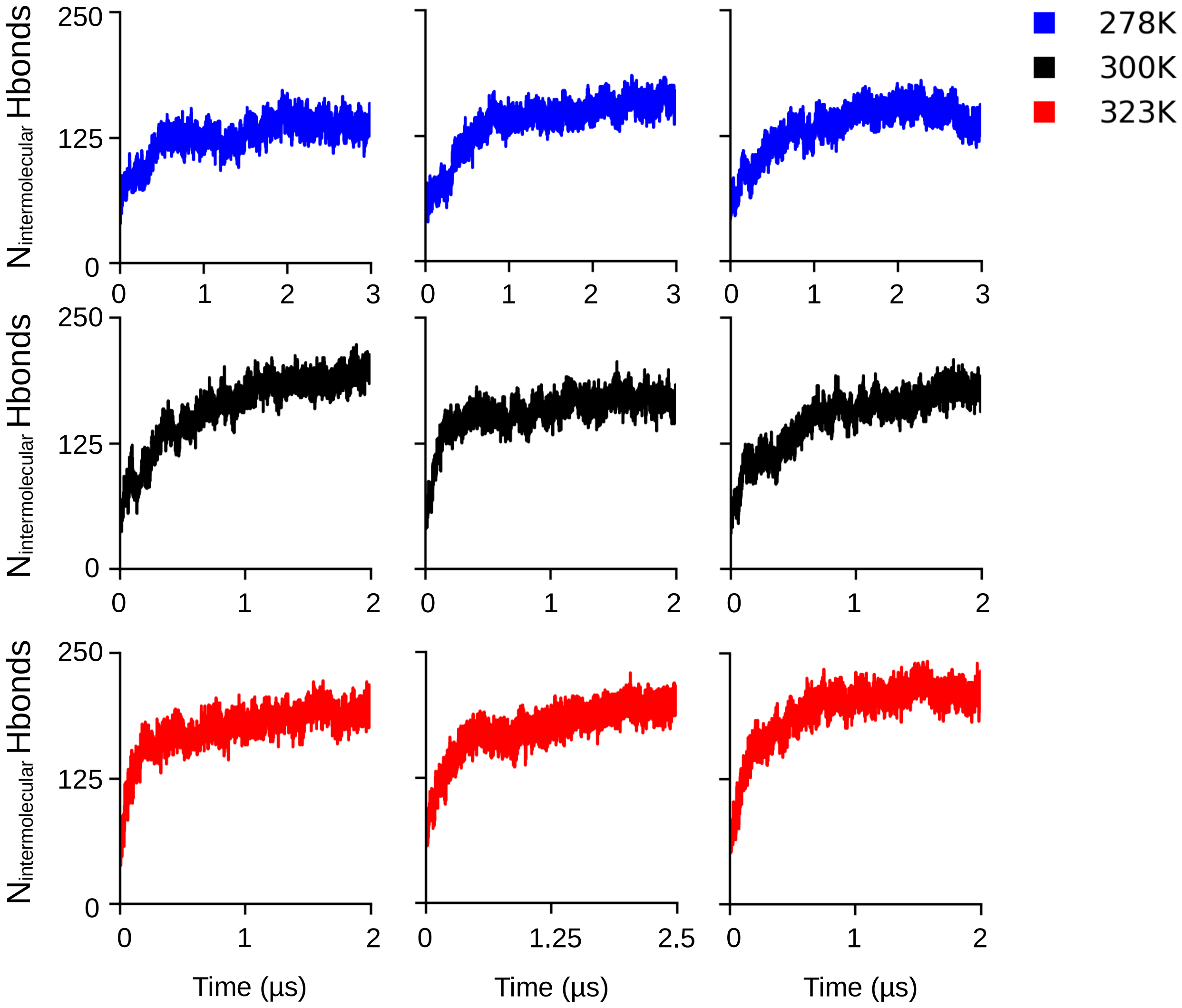


Figure S1: **Number of the interchain hydrogen bonds per acetylglucosamine (GlcAc) unit as a function of simulation time.** A hydrogen bond is considered present if the heavy-atom donor-acceptor distance is below 3.5 Å and the donor-hydrogen-acceptor angle greater than 135°. Note, all 24 chains were used at each temperature.

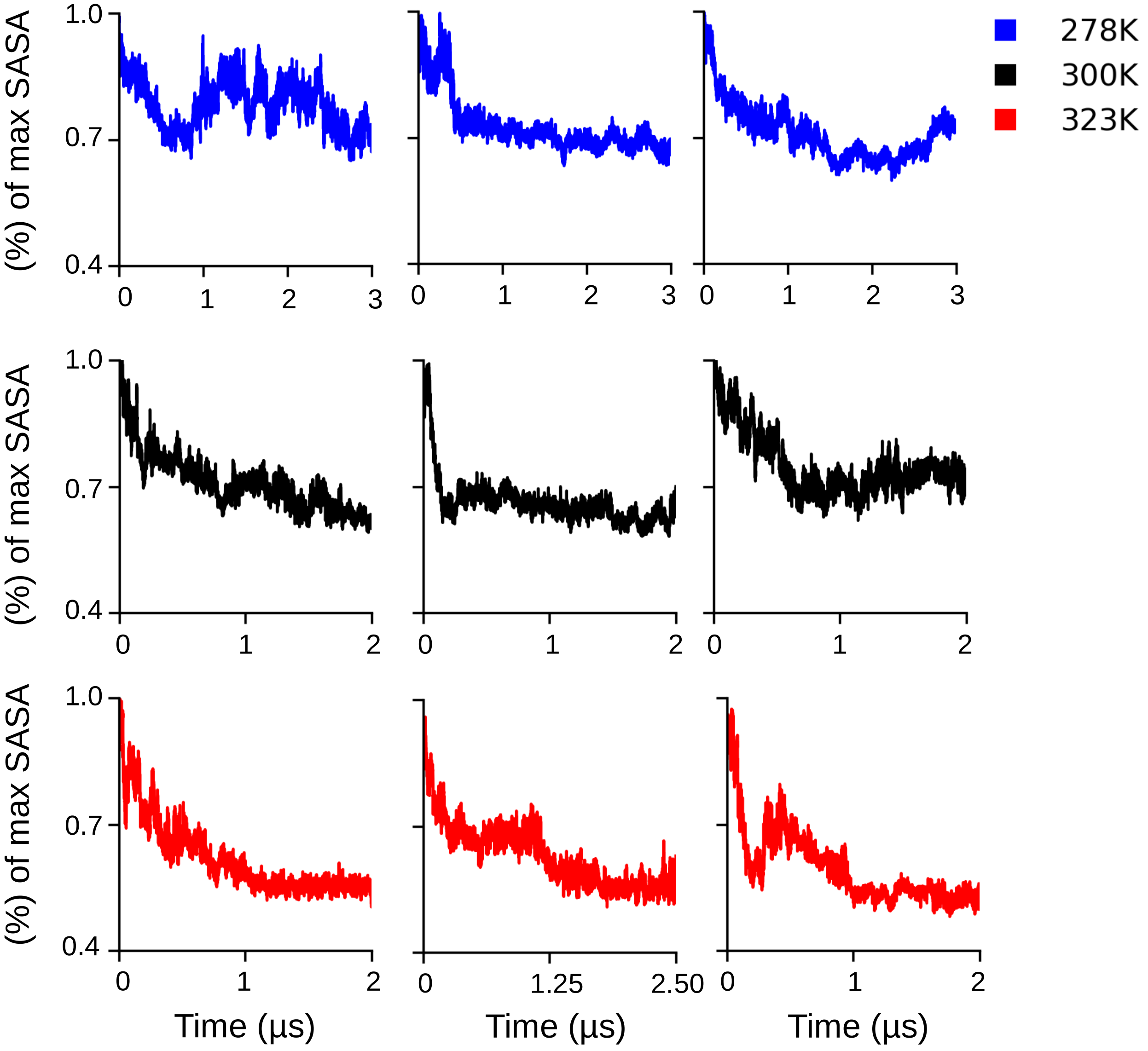


Figure S2: **Fractional solvent accessible surface area (SASA) as a function of simulation time at three temperatures.** SASA is calculated using the LCPO algorithm of Weiser et al. in CPPTRAJ. The fraction of max SASA is calculated by dividing each calculated SASA value by the maximum SASA for that run. Note, all 24 chains were used at each temperature.

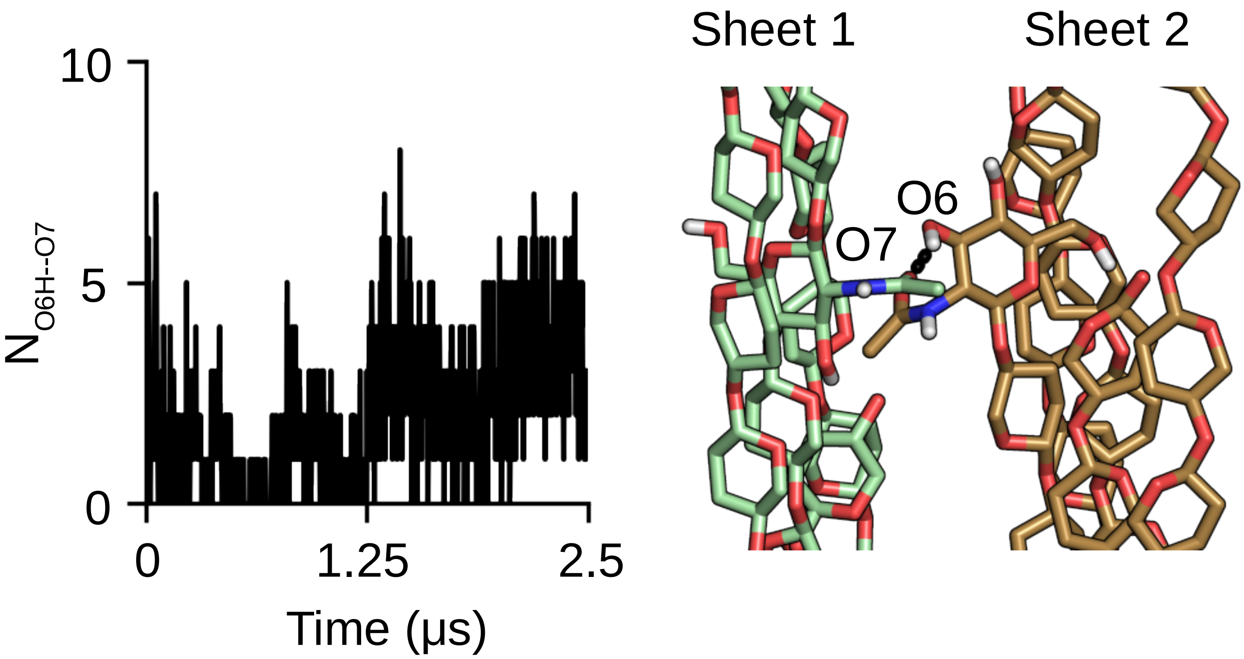


Figure S3: **Number of Intersheet O6H–O7 hydrogen bonding.** Time series of the total number of the most abundant (O6H$\cdots$O7) intersheet hydrogen bonds for the 14-chains that formed a three-sheet nanofibril in the 323 K simulation run 2 (corresponding to Fig. 5a and 5b). A snapshot shows the presence of the O6H$\cdots$O7 hydrogen bond.

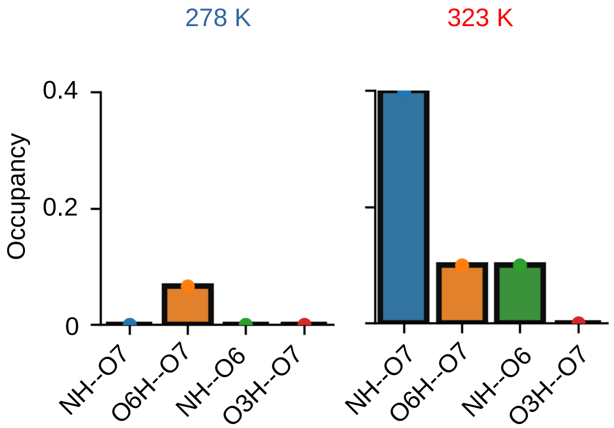


Figure S4: **Specific interchain hbonds at low temperature.** Comparison of the Occupancies of specific interchain hydrogen bonds at 278 K and at 323 K. For 278K Region 1 of run 1 was used for this calculation composed of parallel chains. For 323K region 3 of run 3 was used for the calculation.

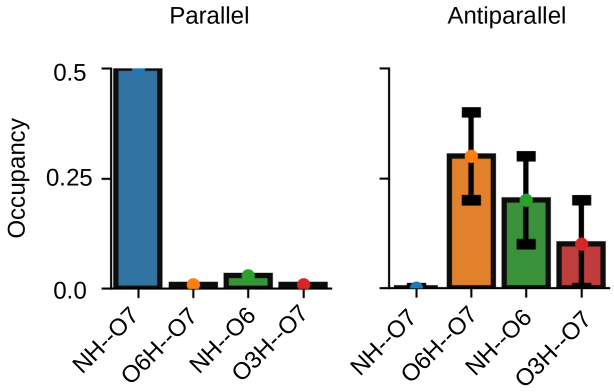


Figure S5: **Specific interchain hbonds at 300K for different chain orientations.** Occupancies of specific interchain hydrogen bonds at 300K for both parallel and antiparallel chains. data taken from run 3. The final 100ns of simulation was used for the calculation
